# Extraction of essential oil, phytochemical screening, and antimicrobial activity study of *Eucalyptus globulus*

**DOI:** 10.1101/2025.08.07.669139

**Authors:** Pragya Neupane, Situ Shrestha Pradhanang

## Abstract

*Eucalyptus globulus* also referred to as Blue Gum is a native plant to Australia where they are cultivated extensively for the extraction of its oil. This oil is of great value in medicinal, therapeutic, and industrial applications. The present study is primarily aimed to the identification of phenolic composition and antimicrobial activity of *Eucalyptus globulus* leaves collected from the Terai region of Nepal. The leaves were partially air-dried, crushed, and subsequently subjected to extraction processes for the preparation of plant extract and essential oil. The GC-MS analysis of essential oil, phytochemical screening, and antimicrobial activity of plant extract were performed. The GC-MS analysis identified a diverse range of compounds, including the major compounds like Eucalyptol(2.52%), Valerianol(5.57%), 10-epi-*γ*-eudesmol(6.93%), Bulnesol(19.97%) and epi-*γ*-eudesmol (17.51%) The antimicrobial efficacy of ethyl acetate (EtOAC), methanol (MeOH), and hexane extracts was evaluated against *Escherichia coli* (ATCC 8739), *Staphylococcus aureus* (ATCC 6538P), and *Candida albicans* (ATCC 2091) using the Mueller-Hinton agar well diffusion method. The hexane extract exhibited significant antibacterial activity, particularly against *E. coli* and *C. albicans*, with inhibition zones of 2.0 cm and 2.1 cm, respectively. In contrast, the MeOH extract showed the least antibacterial activity across all tested strains. The phenolic profiling revealed Bulnesol as the most abundant compound with a concentration of 19.97 units at a retention time of 40.128 minutes, followed by *epi*-*γ*-Eudesmol with 17.51 units. The study underscores the rich phenolic diversity and potent antimicrobial properties of *E. globulus* leaves, highlighting its potential applications in pharmaceuticals and natural product industries.

## Introduction

The global rise in antibiotic-resistant pathogens presents an urgent challenge to public health, creating a critical need for alternative antimicrobial agents. In this context, natural products derived from medicinal plants have emerged as promising sources of bioactive compounds. One such plant, *Eucalyptus globulus labill*., commonly known as the blue gum tree^1^, has shown remarkable potential in traditional and modern medicine. A member of the Myrtaceae family, *E. globulus* is native to Australia but is now widely cultivated across various regions due to its industrial, ecological, and medicinal applications. The plant’s leaves, in particular, are used to extract essential oil that is highly regarded for its broad range of therapeutic properties, including antimicrobial, antioxidant, and anti-inflammatory activities^2–4^.

The essential oil of *Eucalyptus globulus* is predominantly composed of 1,8-cineole (eucalyptol), which can constitute up to 70% of its total content. Alongside 1,8-cineole, the oil contains other significant components such as *α*-pinene, p-cymene, and limonene^2,5^. These compounds have been extensively studied for their biological activities, particularly their antimicrobial effects. Research has demonstrated that the essential oil exhibits potent activity against a range of pathogenic microorganisms, including both Gram-positive and Gram-negative bacteria, as well as fungal strains like *Candida albicans*. This makes *E. globulus* essential oil a valuable candidate for developing natural antimicrobial agents^5,6^.

The safety profile of *Eucalyptus globulus* essential oil is well established, with organizations such as the United States Food and Drug Administration and the Council of Europe recognizing it as safe for use in various consumer products, including pharmaceuticals and cosmetics^7^. Its nontoxic nature, combined with its therapeutic potential, underscores its relevance in current medicinal research. However, despite its established benefits, further exploration into the oil’s phytochemical composition and its specific mechanisms of action remains necessary. Additionally, investigating different extraction methods and solvent systems can help optimize the yield and bioactivity of the essential oil and its related compounds, enhancing its potential applications in combating antibiotic-resistant pathogens.

This research aims to delve deeper into the chemical composition and bioactivity of *Eucalyptus globulus* essential oil and extracts. By employing advanced analytical techniques like Gas Chromatography-Mass Spectrometry (GC-MS)^8^, the study will elucidate the key constituents of the essential oil. Furthermore, different solvent systems, including hexane, ethyl acetate, and methanol, will be used to prepare leaf extracts, offering insights into the efficacy of various extraction methods. The antimicrobial properties of the essential oil and extracts will be tested against a variety of bacterial and fungal strains, including those that exhibit resistance to conventional antibiotics.

Overall, this research will contribute to the growing body of knowledge surrounding natural antimicrobial agents, particularly in the context of antibiotic resistance. By exploring the phytochemical properties and bioactivity of *Eucalyptus globulus*, the study aims to identify compounds that may offer effective alternatives or supplements to traditional antibiotics, addressing a pressing global health challenge.

**Table 1.** Taxonomical classification of *Eucalyptus globulus*.

| Rank | Classification |
| --- | --- |
| Kingdom | Plantae |
| Subkingdom | Tracheobionta |
| Superdivision | Spermatophyta |
| Division | Magnoliophyta (Flowering plants) |
| Class | Magnoliopsida (Dicotyledons) |
| Subclass | Rosidae |
| Order | Myrtales |
| Family | Myrtaceae |
| Genus | *Eucalyptus* |
| Species | *Eucalyptus globulus* Labill. |

*Eucalyptus globulus*, commonly known as Tasmanian blue gum, is a species widely recognized for its rich phytochemical profile, including flavonoids, phenols, and essential oils, which contribute to its antioxidant, antimicrobial, and anti-inflammatory properties^9–11^. Methanolic extracts of *Eucalyptus* leaves have been shown to yield high concentrations of bioactive compounds with potent antioxidant activity^12,13^. Essential oils, particularly those rich in 1,8-cineole, vary significantly across different *Eucalyptus* species. Studies have highlighted the antibacterial efficacy of oils from species such as *E. maideni* and *E. bicostata* against pathogens like *Listeria ivanovii* and *Bacillus cereus*^14^. *Eucalyptus globulus* essential oil (EGEO), predominantly composed of 1,8-cineole, *α*-pinene, and limonene, has demonstrated strong antioxidant, antimicrobial, and insecticidal activities, with significant inhibitory effects against microorganisms such as *Candida tropicalis*^5^. Additionally, phenolic compounds in *E. globulus* fruits, identified through advanced chromatographic techniques, further underscore the plant’s bioactive potential^15^. Recent advancements, such as the synthesis of zinc nanoparticles using *Eucalyptus globulus* extracts, have demonstrated dual antimicrobial and insect-repellent functionalities, offering promising industrial applications^16^. Despite these findings, gaps remain in fully understanding the phytochemical diversity and mechanisms underlying the biological activities of *Eucalyptus globulus*, warranting further research.

## Results

The chemical composition of the essential oil extracted from *Eucalyptus globulus* was analyzed using gas chromatography-mass spectrometry (GC-MS). A total of 25 compounds were identified, with retention times ranging from 9.422 to 46.475 minutes. The major constituents include Bulnesol (19.97%), *epi*-*γ*-Eudesmol (17.51%), and Valerianol (5.79%), indicating these compounds as key contributors to the biological activity of the oil Table 2. The graphical representation of the retention times and compound concentrations Figure 1 shows that Bulnesol and *epi*-*γ*-Eudesmol are the most abundant components, making them dominant in the chemical profile.

**Table 2.**
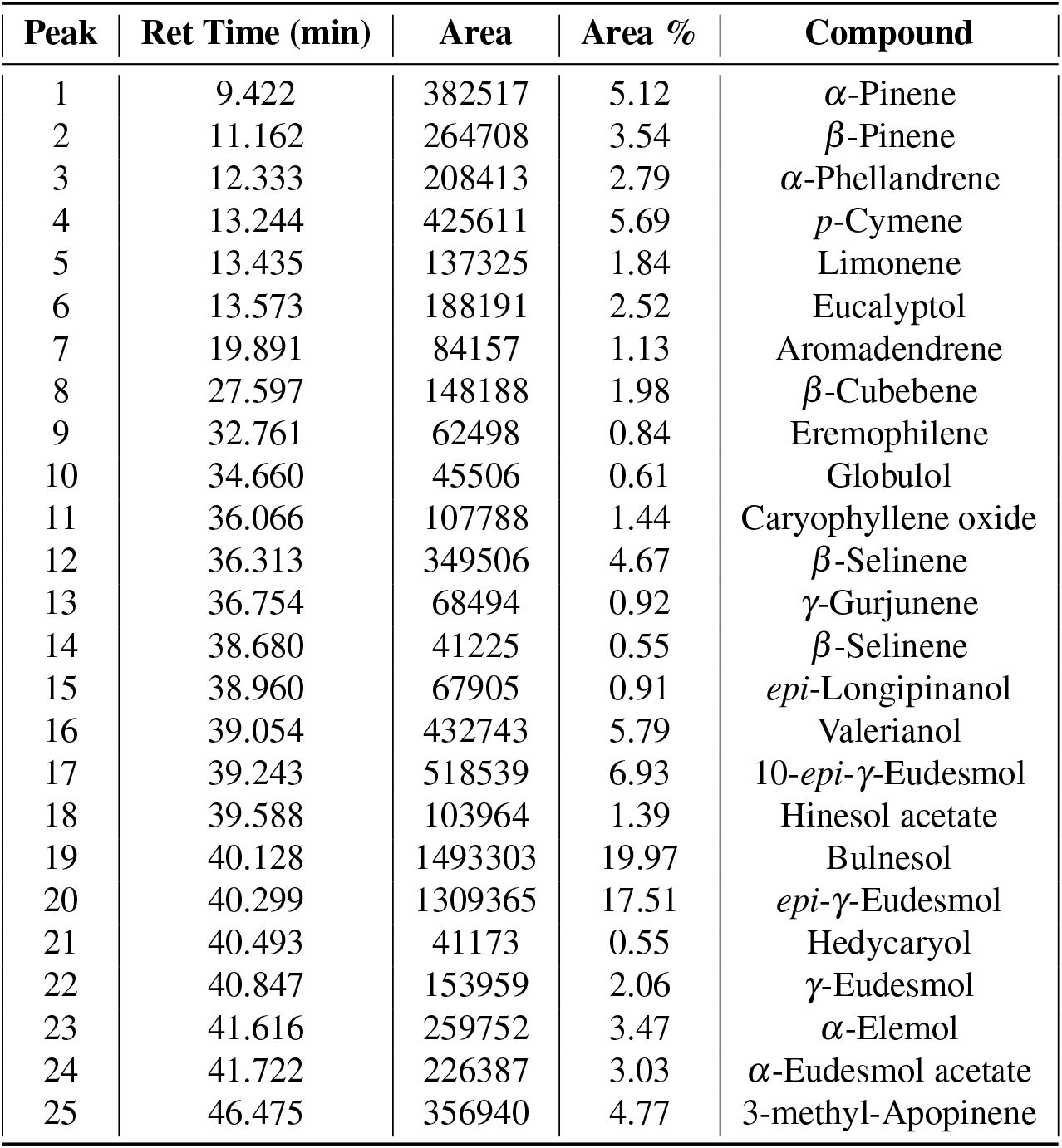
Chemical composition of *Eucalyptus globulus* essential oil identified by GC-MS.

**Figure 1.**
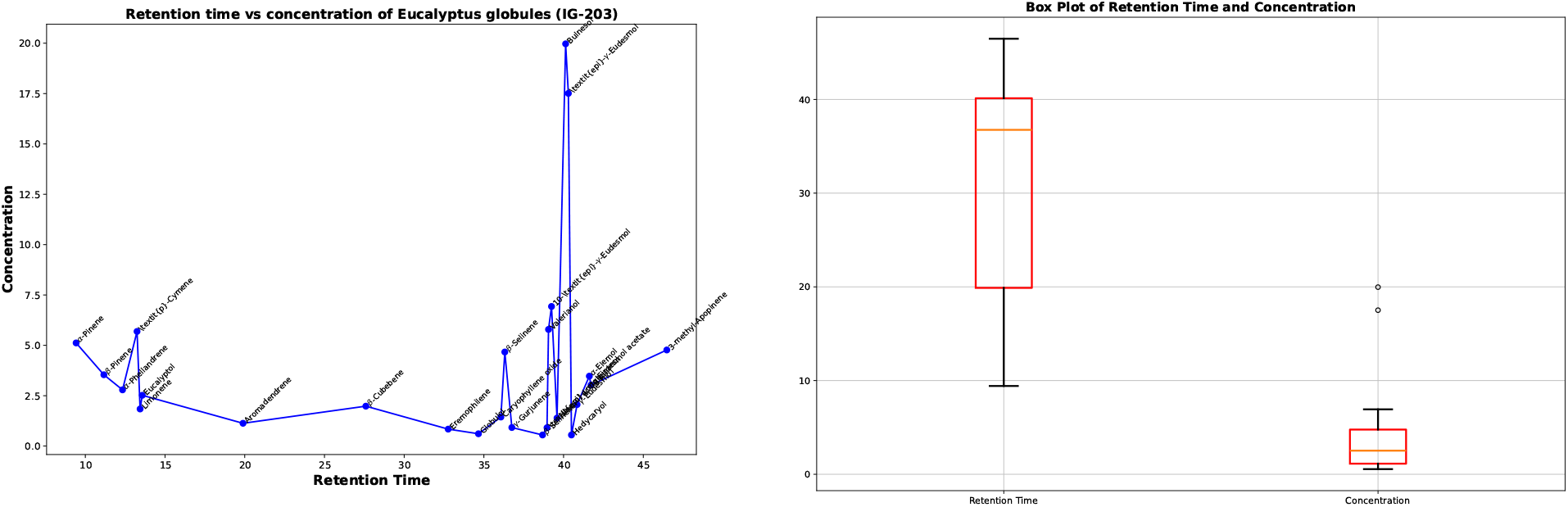
Retention Time vs Concentration of Compounds in *Eucalyptus globulus* essential oil. Box plot of retention time and concentration of compounds in *Eucalyptus globulus* essential oil.

The box plot Figure 1 illustrates the distribution and variability of retention times and concentrations of the compounds. Outliers such as Bulnesol and *epi*-*γ*-Eudesmol emphasize the significant variance in compound concentration, with a median retention time around 36 minutes. The presence of these compounds in high concentrations indicates their relevance to the overall chemical profile of *Eucalyptus globulus* essential oil.

The phytochemical screening of the leaf extracts from *Eucalyptus globulus* confirmed the presence of several key compounds, including flavonoids, polyphenols, glycosides, and terpenoids. The methanol extract showed the highest diversity of phytochemicals, while the hexane extract was particularly rich in terpenoids Table 3.

**Table 3.** Phytochemical screening results of *Eucalyptus globulus* leaf extracts.

| S.N | Constituent | Tests Performed | Hexane | Methanol | Ethyl Acetate |
| --- | --- | --- | --- | --- | --- |
| 1 | Alkaloids | Wagner's test | - | - | - |
| 2 | Flavonoids | Shinoda's test | - | + | + |
| 3 | Polyphenols | Ferric chloride test | - | + | ND |
| 4 | Glycosides | Molisch's test | - | + | ND |
| 5 | Steroids | Lieberman's test | - | - | - |
| 6 | Terpenoids | Salkowski's test | + | - | + |

The antibacterial activity of the extracts was evaluated against three microbial strains: *Escherichia coli* (ATCC 8739), *Staphylococcus aureus* (ATCC 6538P), and *Candida albicans* (ATCC 2091). The hexane extract exhibited the highest antibacterial activity, particularly against *E. coli* and *C. albicans*, as shown in Table 4.

**Table 4.** Antimicrobial activity of *Eucalyptus globulus* extracts against different microbial strains.

| Bacterial strain | Ref. culture | Type | +ve control (c+) | EtOAc ext. | MeOH ext. | Hexane ext. |
| --- | --- | --- | --- | --- | --- | --- |
| *E. coli* | ATCC 8739 | Gram-ve | 2.7 cm | 1.8 cm | 1.3 cm | 2.0 cm |
| *S. aureus* | ATCC 6538P | Gram+ve | 2.2 cm | 1.8 cm | 1.3 cm | 1.6 cm |
| *C. albicans* | ATCC 2091 | Fungi | 2.3 cm | 1.9 cm | 1.3 cm | 2.1 cm |

The hexane extract demonstrated the most promising broad-spectrum antibacterial activity. Compared to the positive control (kanamycin), the hexane extract was particularly effective against *E. coli* and *C. albicans*, with inhibition zones of 2.0 cm and 2.1. cm respectively, closely matching the performance of the positive control.

A comparison with previous studies, such as those by^16^, who synthesized zinc nanoparticles using *Eucalyptus globulus* extracts, showed that the hexane extract in this study exhibited superior antibacterial activity without the need for nanoparticle synthesis. Additionally,^17^ reported similar antimicrobial efficacy in their study of commercial *Eucalyptus globulus* essential oil. These comparisons underscore the potent antimicrobial properties of the natural extracts used in this study, suggesting their potential application in healthcare products and natural antimicrobials.

The analysis of *Eucalyptus globulus* essential oil and leaf extracts revealed a diverse chemical composition with significant antibacterial properties. The hexane extract, in particular, demonstrated broad-spectrum activity, making it a promising candidate for further research into natural antimicrobial agents.

## Discussion

The present study examined the antimicrobial properties of *Eucalyptus globulus* leaf extracts against *Escherichia coli, Staphylococcus aureus*, and *Candida albicans*. The hexane extract exhibited the highest antimicrobial activity, particularly against *E. coli* and *C. albicans*. This potency is likely attributed to the presence of non-polar compounds such as Bulnesol and *epi*-*γ*-Eudesmol, which were identified as the major constituents through GC-MS analysis. These compounds are known for their antimicrobial properties, possibly due to their ability to disrupt microbial cell membranes. The ethyl acetate extract demonstrated moderate antimicrobial activity, while the methanol extract showed the least efficacy. The reduced activity of the methanol extract could be related to its higher concentration of polar compounds, which may not exhibit the same level of antimicrobial effectiveness as non-polar constituents. The chemical composition analysis revealed that the essential oil from *Eucalyptus globulus* is rich in bioactive compounds, including Bulnesol, *epi*-*γ*-Eudesmol, and Eucalyptol (1,8-cineole). Phytochemical screening also confirmed the presence of key compounds such as terpenoids, flavonoids, and phenolic acids, which contribute to the antimicrobial activity of the extracts.

In conclusion, the results of this study highlight the potential of *Eucalyptus globulus* essential oil and extracts as natural antimicrobial agents. The hexane extract, in particular, demonstrated significant activity against bacterial and fungal pathogens, suggesting that non-polar compounds such as Bulnesol and *epi*-*γ*-Eudesmol play a crucial role in its biological efficacy. These findings indicate that *Eucalyptus globulus* could serve as a valuable source for developing natural antimicrobial products. Further studies should focus on isolating and characterizing the individual bioactive compounds from *Eucalyptus globulus* to better understand their specific antimicrobial mechanisms. Additionally, exploring the application of these extracts in various formulations, such as antimicrobial coatings or preservatives, could broaden their potential use in medical, industrial, and agricultural sectors.

## Materials and Methods

The study utilized various apparatus and chemicals essential for the extraction and analysis processes. The apparatus included a Clevenger apparatus^18^ for hydro distillation, a Soxhlet apparatus^19,20^for sequential solvent extraction, a heating mantle, and a condenser^21^. The chemicals employed in the experiments were of analytical grade, including methanol, hexane, ethyl acetate, H_2_SO_4_, HCl, chloroform, Molisch’s reagent, Wagner’s reagent, and DPPH. All chemicals were used without further purification.

The leaves of *Eucalyptus globulus* were collected from the Chandrapur, Rautahat district of Nepal, during March 2024, from an altitude of 110–350 masl, where the species is cultivated extensively for its wood and medicinally valuable leaves. The leaves were washed thoroughly with distilled water to remove impurities, air-dried under shade for two days to preserve phenolic compounds, and hand-crushed Figure 2 before extraction.

**Figure 2.**
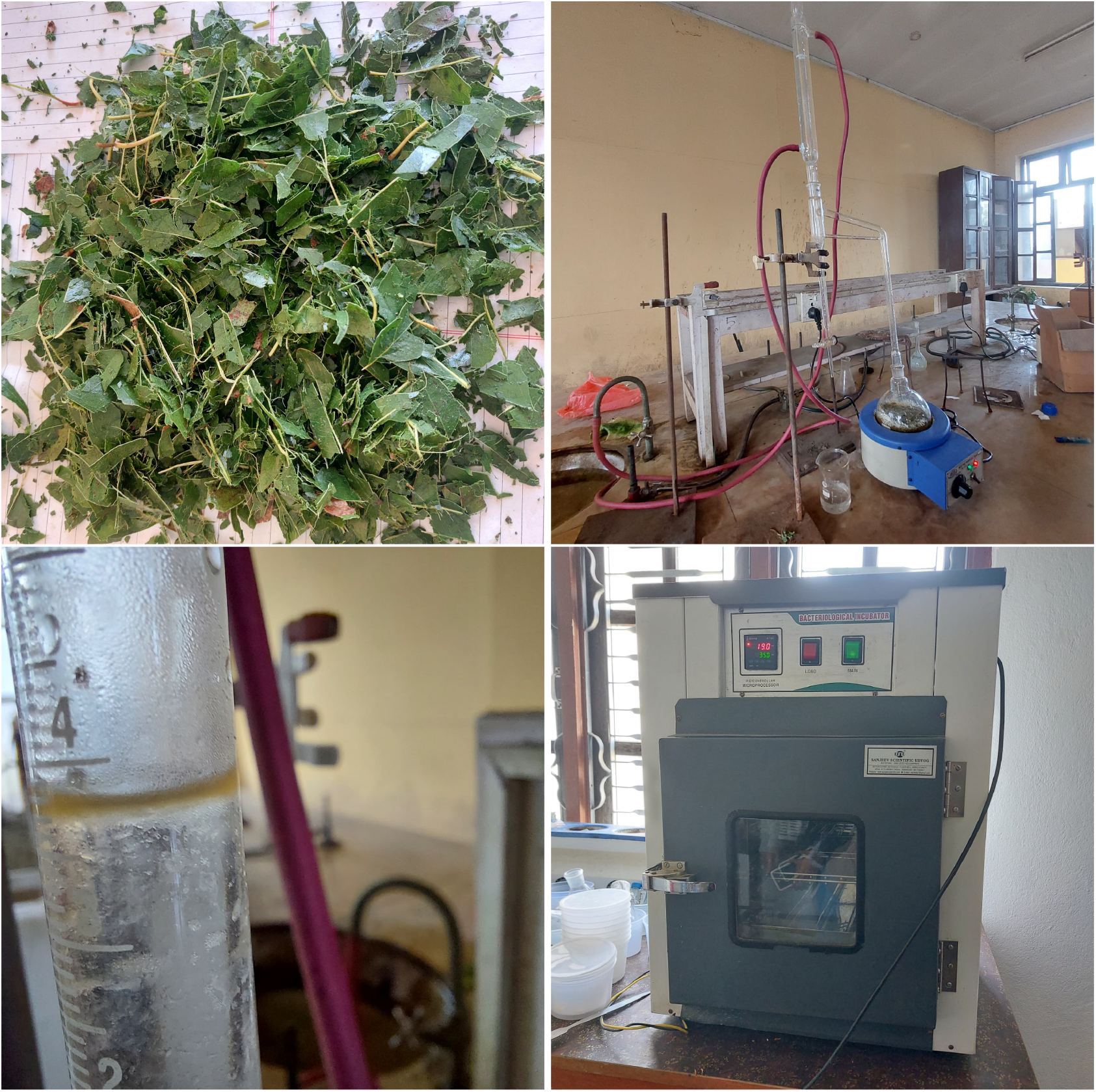
Hand-crushed leaves of *Eucalyptus globulus* prepared for essential oil extraction. Hydrodistillation setup for essential oil extraction using the Clevenger apparatus. A bacteriological incubator was used for the antimicrobial assay.

The essential oil extraction was performed using hydrodistillation with a Clevenger apparatus. A total of 80 g of partially dried leaves were placed in a three-necked round-bottom flask with 200 mL of water. The mixture was heated using a heating mantle, and steam carried the volatile compounds through the condenser. The collected mixture of water and oil was separated, yielding approximately 1 mL of essential oil, rich in compounds like eucalyptol, after two rounds of 90-minute heating cycles Figure 2.

For phytochemical analysis, sequential Soxhlet extraction was carried out using solvents of increasing polarity—hexane, ethyl acetate, and methanol—to extract a broad range of phytochemicals. The air-dried leaves were finely ground and subjected to extraction. The hexane extraction targeted lipophilic compounds, including essential oils and waxes, followed by ethyl acetate extraction for moderately polar compounds like flavonoids. Finally, methanol was used to extract polar compounds such as phenolics and glycosides. The extracts were concentrated by evaporating the solvents and analyzed using gas chromatography-mass spectrometry (GC-MS) to identify the phytochemicals Figure 3.

**Figure 3.**
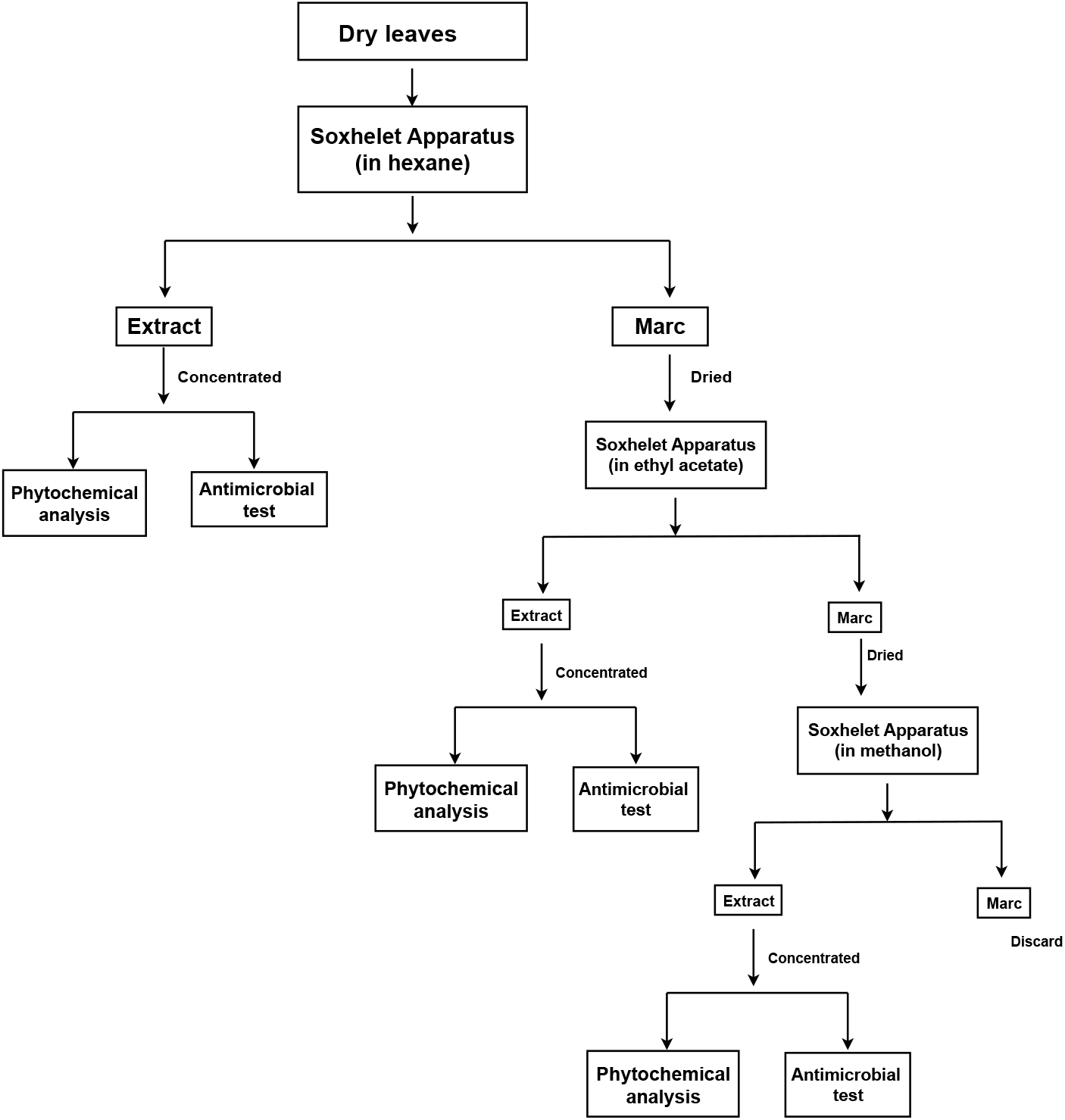
Flowchart depicting the methodology for leaf extract preparation using different solvents.

Phytochemical tests were performed to confirm the presence of various bioactive compounds. Wagner’s test was used to assess the presence of alkaloids, which were found to be absent. Shinoda’s test confirmed the presence of flavonoids, while the Ferric chloride test indicated polyphenols with a bluish-black coloration. Molisch’s test confirmed the presence of glycosides, and Salkowski’s test showed the presence of terpenoids through a reddish-brown interface coloration.

The antimicrobial activity of the extracts was evaluated using the well diffusion method. Mueller-Hinton agar plates were inoculated with bacterial cultures, and wells were loaded with 100 µL of extract solutions at a concentration of 100 mg/mL. Kanamycin (5 mg/mL) served as a positive control. Plates were incubated at 37°C for 24 hours, after which the zones of inhibition were measured Figure 2.

